# Targeted gene correction of *FKRP* by CRISPR/Cas9 restores functional glycosylation of α-dystroglycan in cortical neurons derived from human induced pluripotent stem cells

**DOI:** 10.1101/101352

**Authors:** Beatrice Lana, Jihee Kim, David Ryan, Evangelos Konstantinidis, Sandra Louzada, Beiyuan Fu, Fengtang Yang, Derek L. Stemple, Pentao Liu, Francesco Muntoni, Yung-Yao Lin

## Abstract

Mutations in genes required for functional glycosylation of α-dystroglycan cause a group of congenital muscular dystrophies associated with brain malformations, referred to as dystroglycanopathies. The lack of isogenic, physiology-relevant human cellular models has limited our understanding of the cortical abnormalities in dystroglycanopathies. Here we generate induced pluripotent stem cells (iPSCs) from a severe dystroglycanopathy patient with homozygous mutations in the ribitol-5-phosphate transferase gene, *FKRP*. We carry out targeted gene correction in FKRP-iPSCs using CRISPR/Cas9-mediated genome editing. We characterise the directed differentiation of FKRP- and corrected-iPSCs to neural stem cells, cortical progenitors and cortical neurons. Importantly, we show that targeted gene correction of *FKRP* restores functional glycosylation of α-dystroglycan in iPSC-derived cortical neurons. We independently validate this result by showing targeted gene mutation of *FKRP* disrupts functional glycosylation of α-dystroglycan. This work demonstrates the feasibility of using CRISPR/Cas9-engineered human iPSCs for modelling dystroglycanopathies and provides a foundation for therapeutic development.

**Highlights:**

- Generation of FKRP-iPSCs for modelling cortical abnormalities in dystroglycanopathies
- Precise gene correction by CRISPR/Cas9-mediated genome editing
- Directed differentiation of isogenic control and FKRP-iPSC to cortical neurons
- Functional glycosylation of α-dystroglycan is restored in cortical neurons derived from CRISPR/Cas9-corrected iPSCs
- Targeted gene mutation of FKRP disrupts functional glycosylation of α-dystroglycan in cortical neurons

## Introduction

Post-translational processing of dystroglycan is critical for its function as a cell surface receptor in a variety of fetal and adult tissues. The dystroglycan precursor is cleaved into non-covalently associated α- and β-subunits, forming an integral component of a multiprotein complex. The mature α-dystroglycan is a heavily glycosylated peripheral membrane protein whose molecular weight ranges from 100 to 156 kDa, depending on the tissue-specific glycosylation. In contrast, the β-dystroglycan is a 43 kDa transmembrane protein that links to the actin cytoskeleton via interaction with dystrophin (Barresi and Campbell, 2006). Defective *O*-linked glycosylation of α-dystroglycan is a common pathological hallmark associated with a number of genetic syndromes, encompassing symptoms from muscular dystrophies to ocular defects, cognitive deficits, and cortical malformations (cobblestone lissencephaly) in the central nervous system (CNS). This group of autosomal recessive disorders are commonly referred to as secondary dystroglycanopathies. (Godfrey et al., 2011). Currently there is no cure or effective treatment for dystroglycanopathies. The effort of identifying causative gene mutations in dystroglycanopathies has shed light on a novel mammalian glycosylation pathway (reviewed in Freeze, 2013; Praissman and Wells, 2014; Yoshida-Moriguchi and Campbell, 2015).

To date an increasing number of genes have been implicated in dystroglycanopathies and their products sequentially elaborate the functional glycosylation of α-dystroglycan that is required for binding with extracellular matrix (ECM) ligands, e.g. laminins, perlecan and neurexin (Figure 1A) (Praissman and Wells, 2014; Yoshida-Moriguchi and Campbell, 2015). These include genes involved in the dolichol-phosphate-mannose synthesis: *GMPPB, DPM1, DPM2, DPM3* and *DOLK* (Barone et al., 2012; Carss et al., 2013; Fernandez et al., 2002; Lefeber et al., 2011; Lefeber et al., 2009; Maeda et al., 2000; Ning and Elbein, 2000; Yang et al., 2013); genes required for *O*-mannosylation and subsequent sugar addition: *POMT1, POMT2, POMGNT1, POMGNT2/GTDC2* and *B3GALNT2* (Beltran-Valero de Bernabe et al., 2002; Hiruma et al., 2004; Manya et al., 2004; Manzini et al., 2012; Stevens et al., 2013; van Reeuwijk et al., 2005; Yoshida et al., 2001; Yoshida-Moriguchi et al., 2013) and an *O*-mannose specific kinase gene: *POMK/SGK196* (Jae et al., 2013; Yoshida-Moriguchi et al., 2013). Recent studies have demonstrated the *ISPD* gene encodes a CDP-ribitol pyrophosporylase that generates the reduced nucleotide sugar for the addition of tandem ribitol-5-phosphate to α-dystroglycan by ribitol-5-phosphate transferases, encoded by the *FKTN* and *FKRP* genes (Figure 1A) (Brockington et al., 2001; Gerin et al., 2016; Kanagawa et al., 2016; Kobayashi et al., 1998; Praissman et al., 2016; Riemersma et al., 2015; Roscioli et al., 2012; Willer et al., 2012). Furthermore, *TMEM5* and *B4GAT1/B3GNT1* genes encode enzymes to prime the phospho-ribitol with xylose and then glucuronic acid (Buysse et al., 2013; Praissman et al., 2014; Praissman et al., 2016; Vuillaumier-Barrot et al., 2012; Willer et al., 2014). *LARGE* encodes a bifunctional enzyme to synthesize the subsequent extension of xylose-glucuronic acid disaccharide repeats that function as the binding sites for ECM ligands (Figure 1A) (Inamori et al., 2012; Longman et al., 2003).

**Figure 1.**
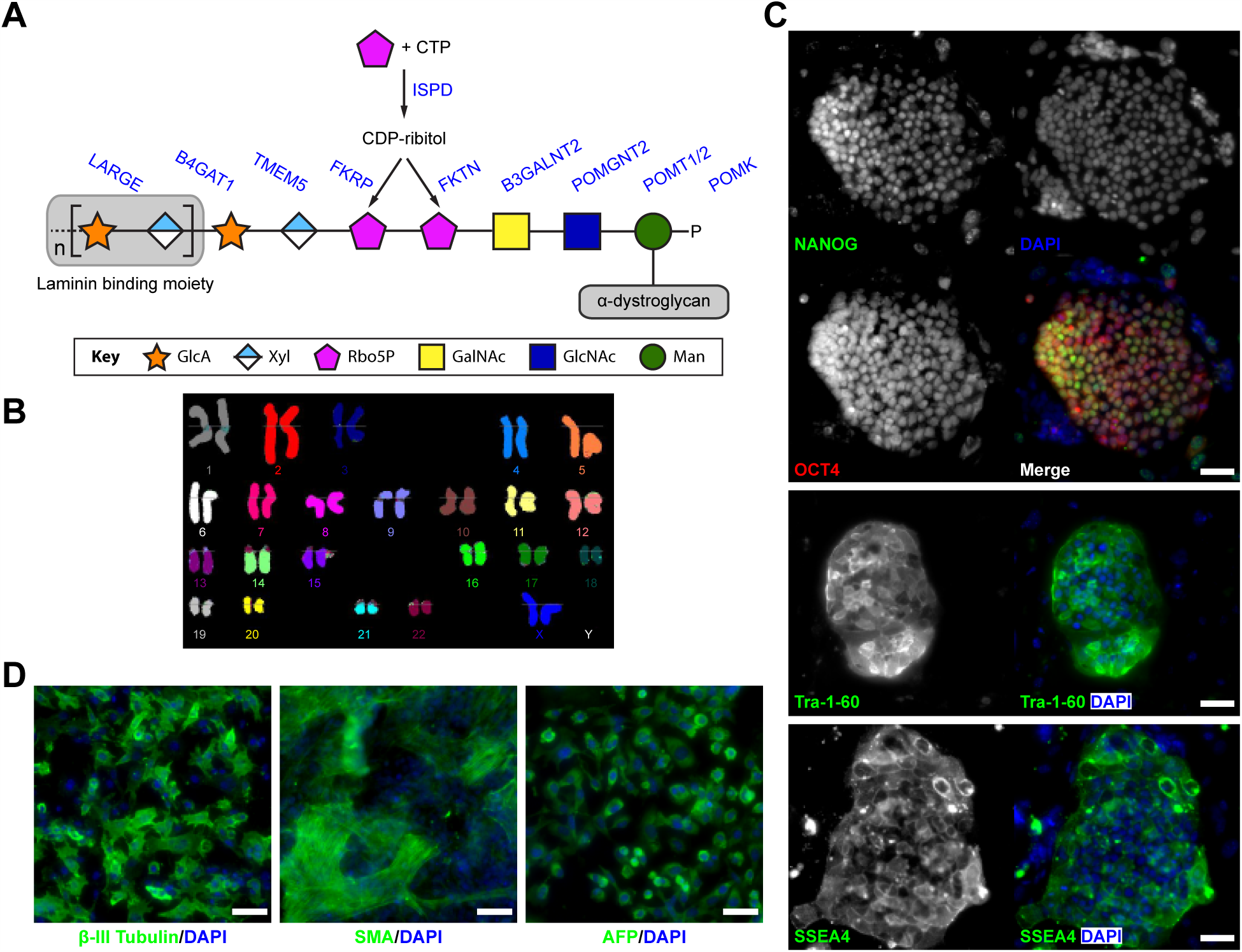
Functional glycosylation of α-dystroglycan and dystroglycanopathy patient-specific iPSCs. (A) Current model of the core M3 functional glycan structure on α-dystroglycan and enzymes involved in its synthesis. ECM ligands bind to the Xyl-GlucA disaccharide repeats. Man, Mannose; GlcNAc, N-Acetylglucosamine; GalNAc, N-Acetylgalactosamine; Rbo5P, ribitol-5-phosphate; Xyl, xylose; GlcA, glucuronic acid. (B) FKRP^A455D^-iPSCs have a normal karyotype. (C) Immunostaining demonstrates FKRP^A455D^-iPSCs express specific pluripotency-associated markers, including NANOG, OCT4, Tra-1-60 and SSEA4. (D) *In vitro* differentiation of FKRP^A455D^-iPSCs to cells representing ectoderm (β-III Tubulin, Tuj1), mesoderm (SMA, smooth muscle actin) and endoderm (α-fectoprotein). Scale bars, 50 μm.

Allelic *FKRP* mutations lead to the widest spectrum of clinical severities, ranging from the mild late-onset limb-girdle muscular dystrophy without neurological deficits (e.g. LGMD2I) to congenital muscular dystrophy with severe CNS abnormalities (e.g. Walker-Warburg syndrome) (Godfrey et al., 2007; Godfrey et al., 2011). Specifically, malformations of cortical development are pathological features at the severe end of the dystroglycanopathy spectrum (Barkovich et al., 2012; Beltran-Valero de Bernabe et al., 2004; Devisme et al., 2012). The absence of physiology-relevant human cellular models has hindered our understanding of mechanisms underlying CNS involvement in dystroglycanopathies and our ability to test potential drug targets in a neural-specific context. In addition, the patient’s neural tissue is not easily accessible and there is no appropriate isogenic control for the patient’s tissue. Induced pluripotent stem cells (iPSCs), which are remarkably similar to embryonic stem cells (ESCs), can be generated from human somatic tissues (Takahashi et al., 2007). Recent studies have demonstrated success in using patient-specific iPSCs to model a variety of neurodevelopment and neurodegenerative disease (An et al., 2012; Ryan et al., 2013; Shi et al., 2012a). These studies suggest that patient-specific iPSC-derived cortical neurons will allow us to study mechanisms underlying CNS involvement in the severe forms of dystroglycanopathies.

In this study we test the hypothesis that patient-specific iPSC-derived cortical neurons can recapitulate pathological hallmarks in dystroglycanopathies. We generate the first human FKRP-iPSCs from a congenital muscular dystrophy patient with severe CNS abnormalities. We carry out CRISPR/Cas9-mediated genome editing (Cong et al., 2013; Mali et al., 2013) and differentiate isogenic pairs of iPSCs to cortical neurons. We show that, for the first time, targeted gene correction of *FKRP* restores α-dystroglycan functional glycosylation in iPSC-derived cortical neurons, whereas targeted gene mutation of *FKRP* disrupts α-dystroglycan glycosylation. Taken together, our isogenic pairs of iPSC-derived cellular models will further elucidate mechanisms underlying the CNS involvement in FKRP-deficient dystroglycanopathies.

## Results

### Generation and characterisation of FKRP-iPSCs derived from a dystroglycanopathy patient with CNS abnormalities

We obtained dermal fibroblasts from a female individual, previously diagnosed congenital muscular dystrophy with CNS abnormalities, including marked cognitive delay, microcephaly, cerebellar cysts and cerebellar dysplasia. Genetic analysis revealed homozygous *FKRP* c.1364C>A (p.A455D) mutations in this affected individual. Both parents were heterozygous carriers of this mutation. To generate patient-specific iPSCs, we reprogrammed the fibroblasts with the *FKRP* c.1364C>A mutations using a six-factor reprogramming technology based on a doxycycline-inducible system (Wang et al., 2011). The initial characterisation of several independent FKRP-iPSC lines by qPCR revealed the gene expression of pluripotency markers, such as *NANOG, OCT4* and *REX1* (Figure S1). Among these patient-specific FKRP-iPSC lines, we focused on the FKRP-iPSC line 1-6, which exhibited the normal karyotype, suggesting these iPSCs are genetically stable (Figure 1B). Immunocytochemistry confirmed the expression of pluripotency markers, such as NANOG, OCT4, Tra-1-60 and SSEA4 in the established FKRP-iPSC line (Figure 1C). In addition, *in vitro* differentiation of the FKRP-iPSCs formed embryoid bodies with cell types representing all three embryonic germ layers, confirmed by immunocytochemistry in specific cell lineages, such as β-III tubulin (Tuj1) for ectoderm, smooth muscle actin (SMA) for mesoderm and α-fetoprotein (AFP) for endoderm (Figure 1D).

### Targeted gene correction of FKRP-iPSCs using CRISPR/Cas9 mediated genome editing

A common issue of patient-specific iPSCs in disease modelling is the lack of appropriate isogenic control cells, causing concerns about the effect of genetic backgrounds on phenotypic variability. To overcome this issue, we applied a precise genome editing strategy to correct the *FKRP* c.1364C>A (p.A455D) mutation using site-specific endonuclease CRISPR/Cas9 stimulated homologous recombination (Cong et al., 2013; Mali et al., 2013), which consists a single guide RNA (sgRNA), the Cas9 nuclease and a donor targeting vector. To avoid potential off-target effects, we used an optimised computational algorithm (http://crispr.mit.edu) to identify appropriate sgRNAs for utilizing the CRISPR/Cas9 to generate DNA double strand breaks (DSB) near the *FKRP* mutation (Figure 2A). This sgRNA has high on-target and low off-target scores (Table S1). To facilitate the screening process for homologous recombination events, we used an approach based on the transposon *piggyBac (PGK-puroΔtk)* selection cassette in the targeting donor vector, which enables both positive and negative selections (Yusa, 2013).

**Figure 2.**
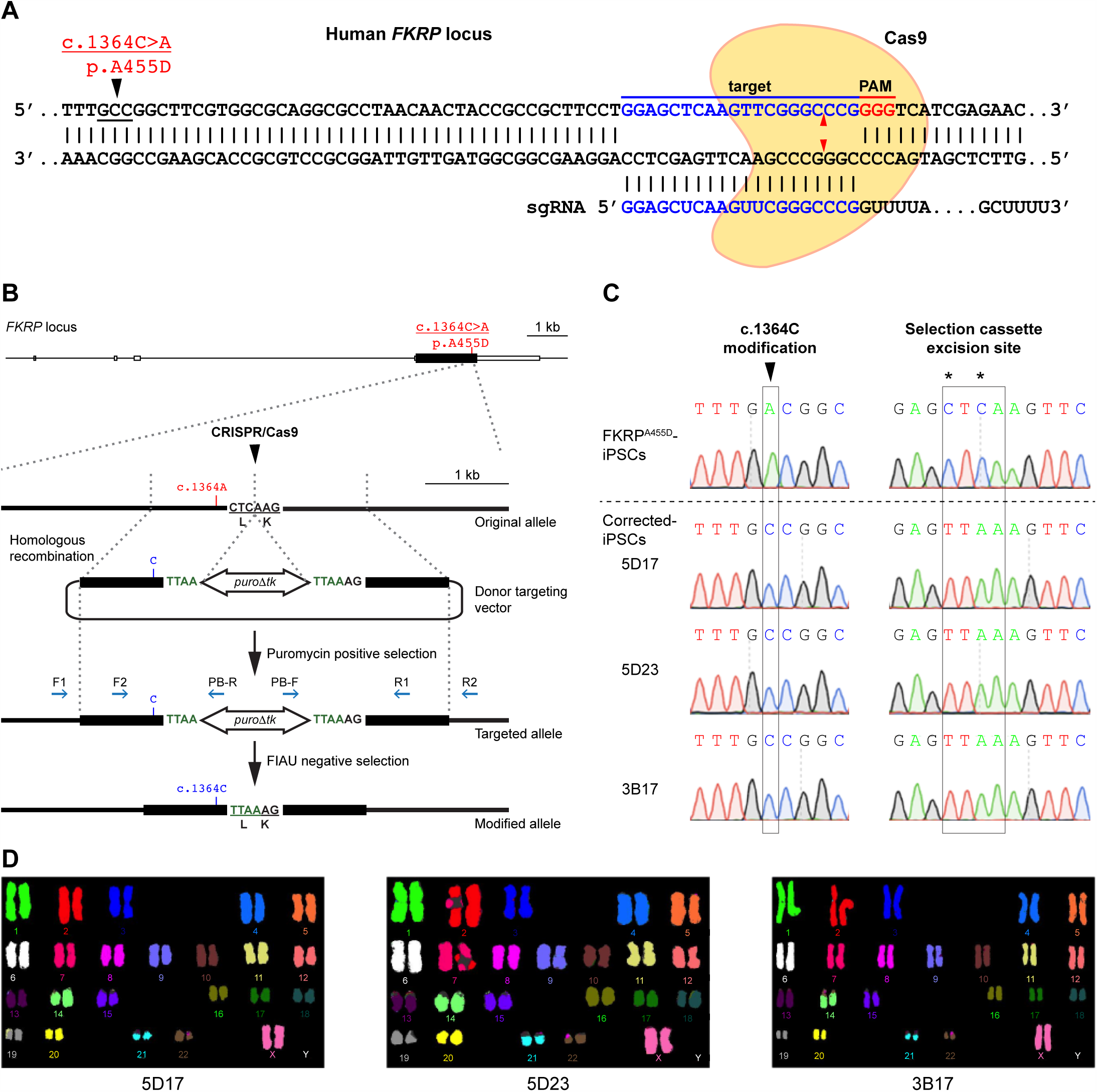
Targeted gene correction of FKRP^A455D^-iPSCs by CRISPR/Cas9-mediated genome editing. (A) Cas9 protein and the specific sgRNA targeting the human *FKRP* locus. The *FKRP* c.1364C>A (p.A455D) mutation is 43 bases upstream of the sgRNA target sequences. Red arrowheads indicate putative cleavage site. PAM, protospacer-adjacent motif. (B) A schematic diagram shows the genome editing strategy based on CRISPR/Cas9-stimulated homologous recombination, followed by positive selection with puromycin and negative selection with FIAU. Homology left and right arms on the targeting donor vector are indicated as black boxes, flanking the *piggyBac (PGK-puroΔtk)* selection cassette, which is under the control of *PGK* promoter. PCR genotyping primers are shown as blue arrows. Note that the TTAA sequences are designed to accommodation the selection cassette excision sites, yet code the same amino acids. (C) Sequence analysis shows precise biallelic correction of the *FKRP* mutation in three independently corrected iPSC clones (5D17, 5D23 and 3B17), compared with their parental FKRP^A455D^-iPSCs. Selection cassette excision sites are identified in the corrected-iPSC lines. (D) CRISPR/Cas9 corrected-iPSC lines show a normal karyotype.

We PCR-amplified two 1-kb fragments flanking the CRISPR/Cas9 target site close to the *FKRP* c.1364C>A allele, which was simultaneously corrected *(FKRP c.1364C).* The two fragments (homology left and right arms) flanking a *piggyBac (PGK-puroΔtk)* selection cassette were assembled together into a targeting donor vector (Figure 2B). DNA sequences at the junctions between *piggyBac (PGK-puroΔtk)* selection cassette and homology arms were modified to accommodate TTAA sequences, yet coding the same amino acids after excision of the selection cassette (Figure 2B). We electroporated the site-specific CRISPR/Cas9 plasmids with the targeting donor vector *(FKRP* c.1364C) into the iPSCs. By positive selection with puromycin, iPSCs with an integrated donor vector formed puromycin-resistant clones, which were picked for rapid PCR genotyping (Figure S2). We identified 3 homozygously and 27 heterozygously targeted independent clones (Table 1), which were confirmed by sequencing (data not shown). To excise the *piggyBac (PGK-puroΔtk)* selection cassette, 2 homozygously targeted *FKRP* (c.1364C)-iPSC clones (5.19 and 3.16) were electroporated with *piggyBac* transposase expression plasmid, followed by negative selection in culture media containing 1-(2-Deoxy-2-fluoro-β-D-arabinofuranosyl)-5-iodo-2,4(1H,3H)-Pyrimidinedione (FIAU), a thymidine analogue that would be processed to toxic metabolites in the presence of *piggyBac (PGK-puroΔtk).* PCR genotyping identified 11 biallelicly corrected-iPSC clones that had the selection cassette completely excised without re-integration (Figure S3 and Table 1). The biallelicly corrected-iPSC clones (5D17, 5D23 and 3B17) were sequenced to confirm a clean correction of the *FKRP* mutation and the engineered selection cassette excision site (Figure 2C). The biallelicly corrected iPSC lines retained normal karyotypes (Figure 2D) and pluripotency (data not shown). In addition, we sequenced the top 5 potential off-target sites (Table S1) and confirmed that no mutations were introduced during genome editing (data not shown).

**Table 1.**
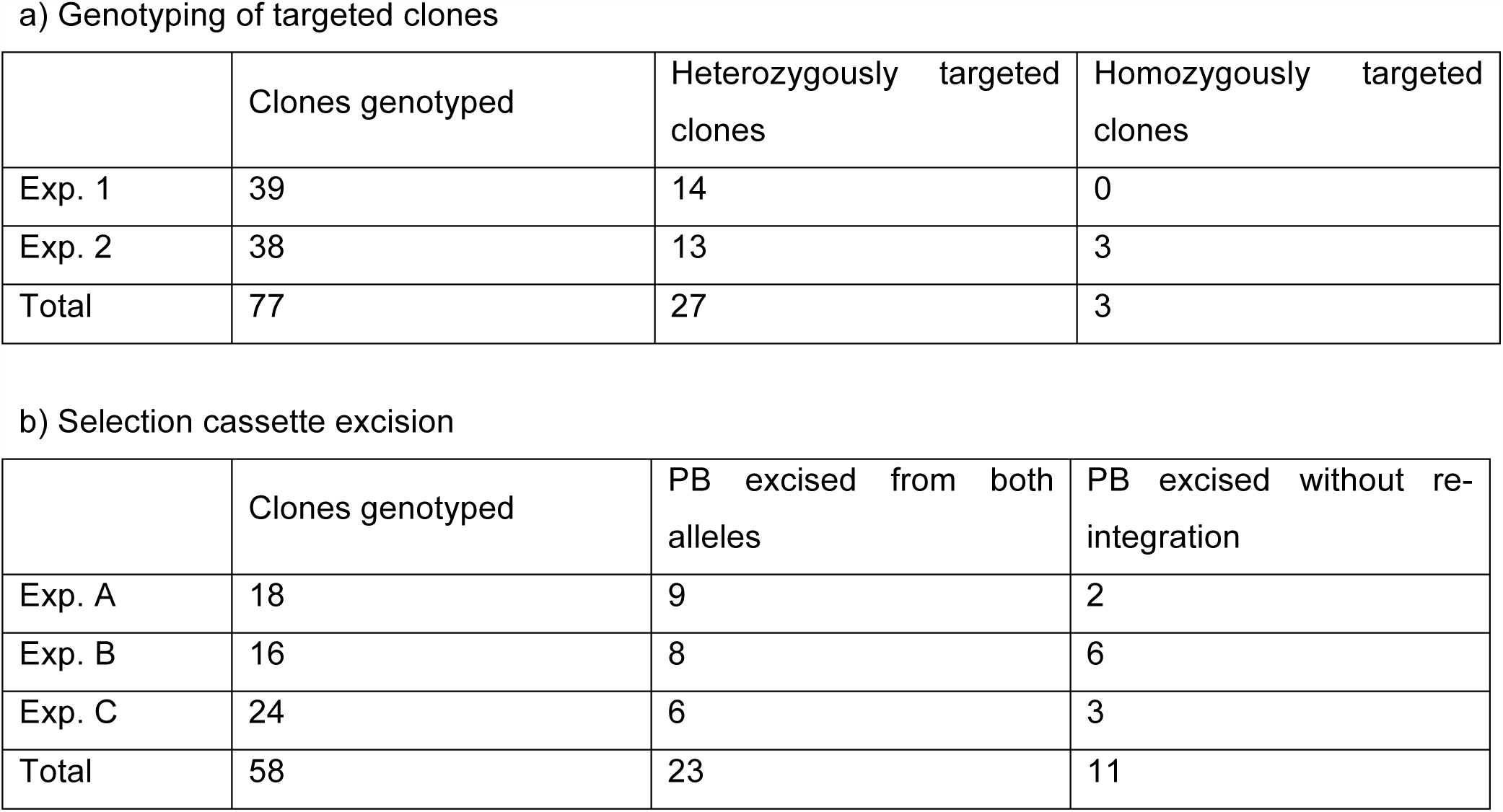
Targeted gene correction by CRISPR/Cas9

### Generation of neural stem cells and cortical neurons from FKRP- and CRISPR/Cas9 corrected-iPSCs

To investigate the potential of iPSCs for modelling neural pathogenesis in dystroglycanopathies, we used a serum-free neural induction medium to derive primitive neural stem cells (NSCs) from iPSCs (Yan et al., 2013). Immunocytochemistry confirmed that NSCs derived from FKRP- and CRISPR/Cas9 corrected-iPSC lines express classic NSC markers, including SOX1, SOX2 and nestin (Figure 3A,B). In terms of the efficiency of neural induction, we did not observe discernable difference between NSCs derived from FKRP- and three corrected-iPSC lines, 5D17, 5D23 and 3B17 (Figure 3C,D). Quantification of SOX1 and SOX2 positive NSCs showed >99% efficiency from both FKRP- and three corrected iPSC lines (Figure 3C,D).

**Figure 3.**
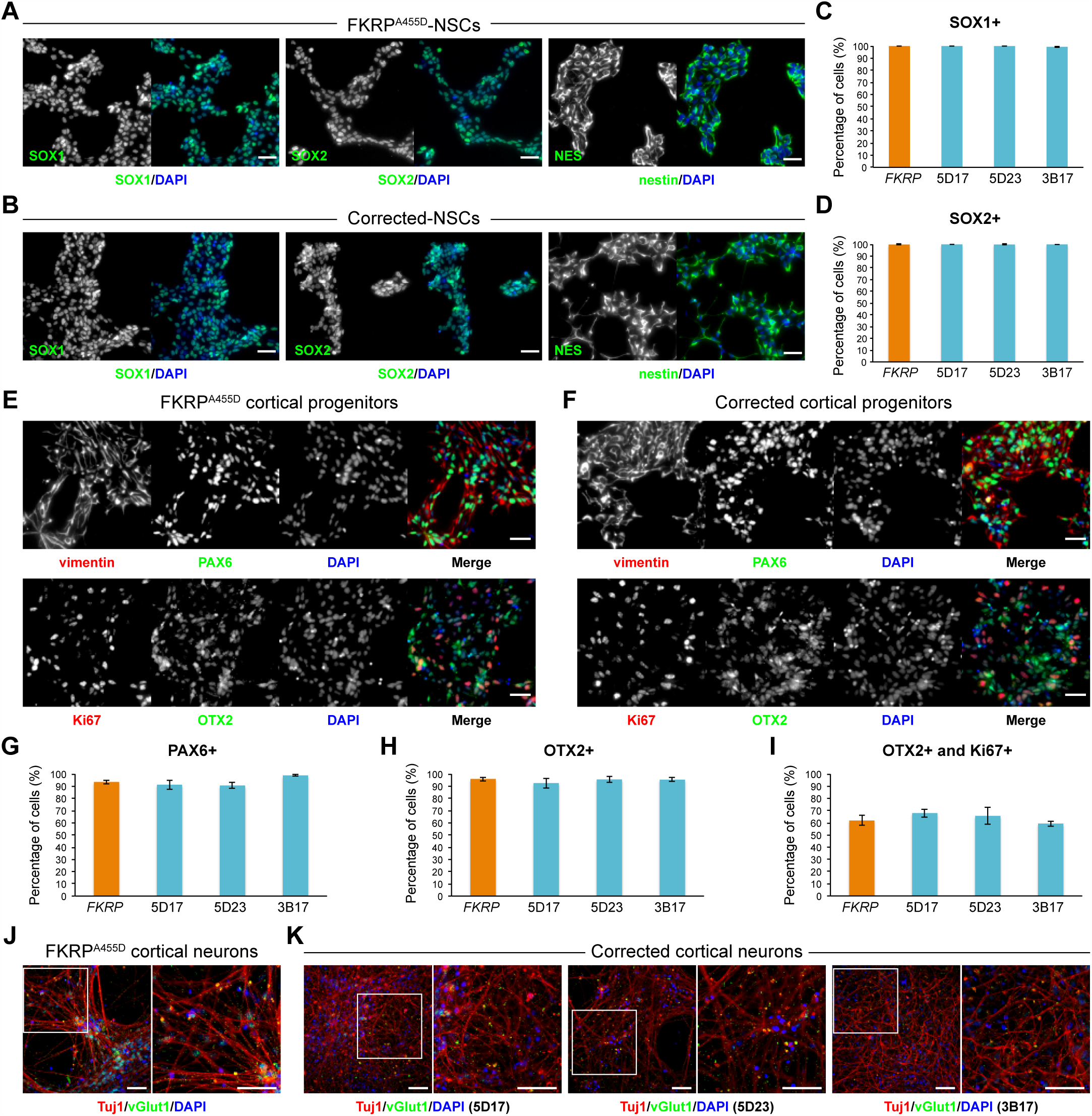
Characterisation of NSCs and cortical neurons derived from FKRP- and CRISPR/Cas9 corrected-iPSCs. (A and B) Representative images of NSCs derived from FKRP- and corrected-iPSC lines expressing SOX1, SOX2 and nestin. (C and D) Quantification of percentage of SOX1+ (C) and SOX2+ (D) cells in culture. The efficiency of neural induction is more than 99% in FKRP- and corrected-iPSC (5D17, 5D23 and 3B17) lines. (E and F) FKRP- and corrected NSC lines can be further differentiated to cortical neural progenitor cells, expressing PAX6, OTX2 and vimentin. (G-I) Quantification of percentage of PAX6+ (G) and OTX2+ (H) cells in culture. About 91-98% of cells derived from *FKRP*, 5D17, 5D23 and 3B17 NSC lines express PAX6 (G). About 93-96% of cells derived from *FKRP*, 5D17, 5D23 and 3B17 NSC lines express OTX2 (H). Of the OTX2+ population, about 60-67% cells are also Ki67+ cycling progenitors (I). (J and K) Glutamatergic projection neurons derived from FKRP and corrected (5D17, 5D23 and 3B17) progenitor cells. The vast majority of neurons contain vGlut1+ punctae in their neurites (labelled by Tuj1). Right panels are enlarged images from the insets of left panels. Scale bars, 50 μm.

Subsequently, we directed the differentiation of FKRP- and corrected-NSC lines towards cortical projection neurons using a well-established protocol, which recapitulates important stages in human cortical development (Shi et al., 2012b). After switching to the Neural Maintenance Medium for one week, we confirmed the identity of FKRP- and corrected-iPSC derived cortical stem and progenitor cells, which express classic cortical stem cell markers, PAX6, OTX2 and vimentin (Figure 3E,F), and the proliferating cells are Ki67 positive (Figure 3E,F). The efficiency of cortical induction between FKRP and three lines of corrected cortical progenitors are very similar. The PAX6+ cells in culture are about 91-98% and OTX2+ cells are about 93-96% (Figure 3G,H). We found 60-67% of OTX2+ cells are also Ki67+ cycling progenitors (Figure 3I). After three weeks in Neural Maintenance Medium, FKRP and corrected progenitor-derived cells showed punctate staining of vesicular glutamate transporter 1 (vGlut1) in their neurites labelled by neuron-specific tubulin, Tuj1 (Figure 3J,K), confirming the generation of glutamatergic projection neurons during cortical neurogenesis in culture.

### Functional glycosylation of α-dystroglycan is restored in cortical neurons derived from CRISPR/Cas9 corrected-iPSCs

Next, we investigated whether we could detect functional glycosylation of α-dystroglycan in cortical neurons derived from our corrected-iPSC lines. To do this, we performed immunoblot using the IIH6 antibody, which recognises the laminin-binding glyco-epitope on α-dystroglycan (Michele et al., 2002). Wildtype mouse muscle and brain lysates were used as positive controls to show differential glycosylation of α-dystroglycan in a tissue-specific manner. In addition, we included lysate of cortical neurons from a non-isogenic, wildtype iPSC20 line (hereinafter called WT-iPSCs) (Wang et al., 2011). We showed that the molecular weight of glycosylated α-dystroglycan in WT-iPSC derived cortical neurons is similar to that in mouse brain lysate (˜120 kDa) and less than that in muscle lysate (˜156 kDa) (Figure 4A), consistent with previously reported tissue-specific glycosylation of α-dystroglycan (Barresi and Campbell, 2006). Furthermore, IIH6 reactivity on immunoblot was almost not detected in cortical neurons derived from FKRP-iPSCs (Figure 4A and B). In contrast, cortical neurons derived from CRISPR/Cas9 corrected-iPSC lines clearly showed IIH6 reactivity, indicating restored glycosylation of α-dystroglycan (Figure 4B).

**Figure 4.**
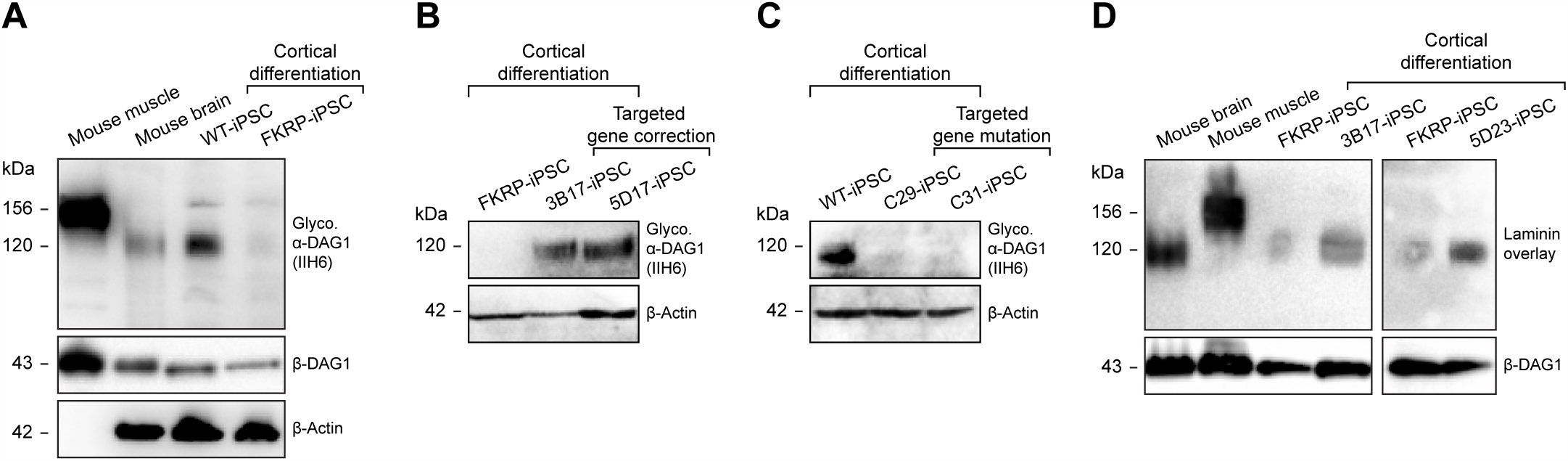
Functional glycosylation of α-dystroglycan in cortical neurons derived from CRISPR/Cas9-engineered iPSCs. (A) Molecular weight of glycosylated α-dystroglycan (IIH6 epitope) in WT-iPSC derived cortical neurons is similar to that in the mouse brain (˜120 kD) and lower than that in the mouse muscle (˜156kD). IIH6 reactivity was almost not detected in cortical neurons derived from FKRP-iPSCs. Note that the β-Actin antibody does not cross-react with the muscle sample. (B) Targeted gene correction of *FKRP* restores IIH6 reactivity in iPSC-derived cortical neurons. (C) Targeted gene mutation of *FKRP* disrupts IIH6 reactivity in iPSC-derived cortical neurons. (D) Targeted gene correction restores laminin-binding activity in iPSC-derived cortical neurons.

Following the detection of glycosylated α-dystroglycan in corrected cortical neurons, we investigated whether the laminin-binding activity is associated with the IIH6 glyco-epitope on α-dystroglycan. We confirmed that laminin-binding is disrupted in cortical neurons derived from FKRP-iPSCs (Figure 4D), whereas cortical neurons derived from corrected-iPSC lines showed strong laminin-binding activities (Figure 4D). Together, these results demonstrated a restoration of functional glycosylation of α-dystroglycan in CRISPR/Cas9-corrected corrected cortical neurons.

### Targeted gene mutation of *FKRP* disrupts functional glycosylation of α-dystroglycan in cortical neurons derived from iPSCs

To independently validate the results from our FKRP- and corrected-iPSC derived cortical neurons, we used the same genome editing strategy (Figure S4A) to knock-in the *FKRP* c.1364C>A (p.A455D) mutation into the WT-iPSCs (Wang et al., 2011). We constructed a donor targeting vector carrying the *FKRP* mutation for CRISPR/Cas9-mediated homologous recombination. We electroporated the site-specific CRISPR/Cas9 plasmids with the targeting donor vector *(FKRP* c.1364C>A) into the WT-iPSCs (Figure S4B). Using puromycin positive selection and PCR genotyping (Figure S2), we identified 2 homozygously targeted clones (a15 and c43), which were expanded for selection cassette excision (Table 2). Following FIAU negative selection and PCR genotyping (Figure S3), we identified 2 biallelicly *FKRP* mutated-iPSC clones (C29 and C31) that have the selection cassette completely excised without re-integration (Table 2). The *FKRP* mutated-iPSC clones (C29 and C31) were sequenced to confirm the biallelic knock-in of *FKRP* c.1364C>A (p.A455D) mutations and the engineered selection cassette excision site (Figure S4C). The wildtype- and *FKRP* mutated-iPSC lines were differentiated to NSCs (Figure S5) and subsequently cortical neurons for functional analysis. Compared with cortical neurons derived from WT-iPSCs, targeted gene mutation of *FKRP* disrupted the IIH6 reactivity (Figure 4A and C) in cortical neurons derived from *FKRP* mutated-iPSC lines, indicating loss of functional glycosylation of α-dystroglycan. Together, our results demonstrated that target gene mutation of *FKRP* by CRISPR/Cas9-mediated genome editing could recapitulate pathological hallmarks of dystroglycanopathy in cortical neurons.

**Table 2.**
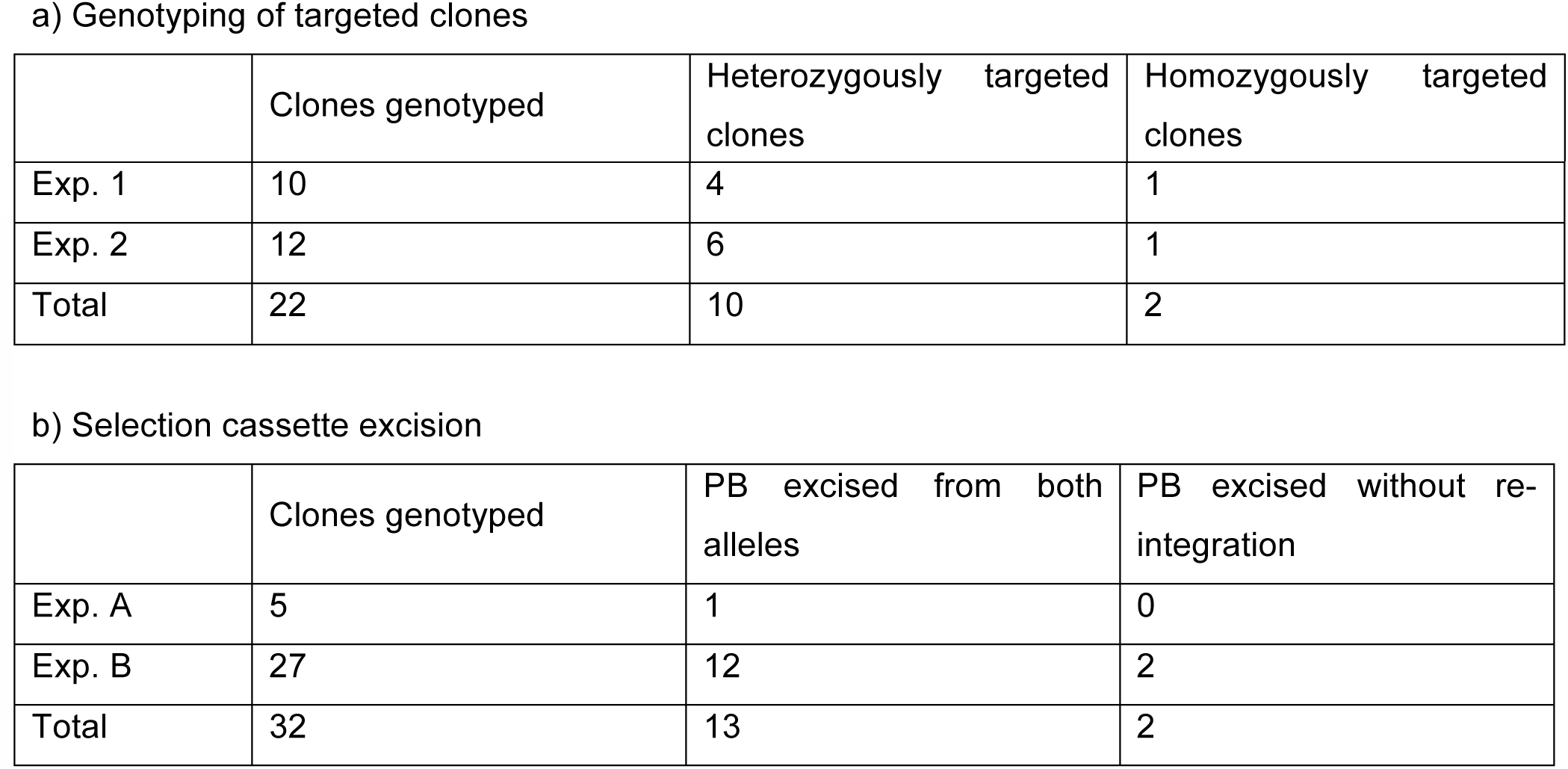
Targeted gene mutation by CRISPR/Cas9

a) Genotyping of targeted clones
|  | Clones genotyped | Heterozygously targeted clones | Homozygously targeted clones |
| --- | --- | --- | --- |
| Exp. 1 | 10 | 4 | 1 |
| Exp. 2 | 12 | 6 | 1 |
| Total | 22 | 10 | 2 |

b) Selection cassette excision
|  | Clones genotyped | PB excised from both alleles | PB excised without re-integration |
| --- | --- | --- | --- |
| Exp. A | 5 | 1 | 0 |
| Exp. B | 27 | 12 | 2 |
| Total | 32 | 13 | 2 |

## Discussion

The combination of patient-specific iPSCs and CRISPR/Cas9-mediated genome editing has enormous potential for disease modelling, as well as for drug discovery and development. We report here that CRISPR/Cas9-induced homologous recombination together with the *piggyBac* positive/negative selection cassette is a powerful and versatile strategy, which allows precise modification of the mammalian genome at a single base-pair level without leaving footprints. For the first time, we demonstrate a precise genome editing at the *FKRP* locus in human iPSCs. Using our genome editing strategy, we were able to achieve 7-10% bialleic targeting efficiency at the *FKRP* locus (Table 1) and no mutations were observed at the top 5 predicted off-target sites (data not shown). According to Leiden Open Variation Database (http://www.lovd.nl/), at least 95% reported *FKRP* DNA variants are missense mutations (969 out of 1018 cases). Homozygous null *FKRP* mutations are exceptionally rare and a single case of Walker-Warburg syndrome with a mutation affecting the translational start site was reported (Van Reeuwijk et al., 2010). Most patients carry heterozygous or homozygous *FKRP* mutations or, less frequently, a combination of a nonsense or frameshift mutation with a missense mutation (5 out of 38 patients in our research cohort). Among these mutations, *FKRP* c.826C>A (L276I) mutation is one of the most common variants and causes a mild form of dystroglycanopathy (LGMD2I). Our genome editing strategy can be applied to precisely correct these *FKRP* missense mutations and by implication small insertion/deletion mutations (INDELs), as well as similar type of mutations in other dystroglycanopathy genes. Recent advances in the delivery of Cas9-sgRNA ribonucleoproteins (Cas9 RNPs) (Liu et al., 2015), may further improve the efficiency of biallelic gene targeting. The advantage of using Cas9 RNPs for genome editing is the rapid Cas9 cleavage action and protein turnover in the cells within 24 h of delivery, which increases the gene targeting efficiency and reduces the off-target mutation rate that is critical for future therapeutic application.

The process of cellular reprogramming from fibroblasts to iPSCs followed by directed differentiation from iPSCs to NSCs and subsequently to cortical neurons involves dramatic changes of gene expression profile of a cell. The functional glycosylation of α-dystroglycan is a consequence of a complex biosynthetic pathway orchestrated by at least 17 known enzymes to form the ECM ligand-binding moiety (Yoshida-Moriguchi and Campbell, 2015). Our results indicate that genes involved in functional glycosylation of α-dystroglycan are expressed during *in vitro* corticogenesis. Interestingly, the molecular weight of α-dystroglycan varies due to tissue-specific O-glycosylation (Barresi and Campbell, 2006). Consistent with previous studies, we show that the molecular weight of glycosylated α-dystroglycan in cortical neurons derived from human iPSCs is similar to that in the mouse brain and less than that in mouse muscle (Figure 4). Importantly, we demonstrate that targeted gene correction of *FKRP* restores functional glycosylation of α-dystroglycan in iPSC-derived neurons, whereas targeted gene mutation of *FKRP* disrupts functional glycosylation of α-dystroglycan (Figure 4). Our findings have significant implications in using human iPSCs for modelling dystroglycanopathies because the *in vitro* cellular model can recapitulate the *in vivo* pathological hallmarks of the target tissues.

Mutations in *FKRP* and other known causative genes together account for approximately 50-60% of severe forms of dystroglycanopathies (Roscioli et al., 2012; Vuillaumier-Barrot et al., 2012). Previously a zebrafish based study showed cooperative interaction between *ISPD, FKTN* and *FKRP* (Roscioli et al., 2012). Recent studies demonstrated the enzymatic function of ISPD, FKTN and FKRP, which synergistically add tandem ribito-5-phosphate onto α-dystroglycan (Gerin et al., 2016; Kanagawa et al., 2016). Despite these advances, the effects of *FKRP* mutations and defective glycosylation of α-dystroglycan in a disease context still remain poorly understood. Our isogenic pairs of control- and FKRP-iPSC derived cortical neurons will further elucidate molecular mechanisms underlying cortical development in health and disease using a systems-based approach, similar to transcriptome analysis of *in vitro* corticogenesis from human ESCs (van de Leemput et al., 2014). Importantly, this work will not only elucidate the role of glycosylated dystroglycan in the brain, but will also have significant impacts on studying its role in other tissues. Apart from muscle degeneration, dilated cardiomyopathy is a frequent associated feature of FKRP-deficient dystroglycanopathy (Mercuri and Muntoni, 2013). The isogenic pairs of human iPSCs generated in this study have enormous potential in many other research avenues. In addition to neural differentiation, the isogenic pairs of control- and FKRP-iPSC lines can be used for myogenic and cardiac differentiation with established protocols (Bellin et al., 2013; Chal et al., 2016) for studying pathological mechanisms underlying muscle degeneration and dilated cardiomyopathy in muscular dystrophy patients. In addition, one can envisage the use of progenitor cells derived from CRISPR/Cas9-corrected iPSCs for autologous cell replacement therapies in the future. The advantage of using autologous, CRISPR/Cas9-corrected progenitor cells is the reduced risk of immunological rejection and need for immunosuppression.

In conclusion, we have established two isogenic pairs of human iPSC models for a severe form of FKRP-deficient dystroglycanopathy. A significant impact of this study is that other human iPSC-based dystroglycanopathy models will be generated. Collectively, these iPSC models will provide powerful *in vitro* platforms that can be exploited for setting up high-content drug screens using quantifiable phenotypes as readouts. Importantly, while more than 90% of drugs fail during development (DiMasi et al., 1991), our isogenic pairs of human iPSC lines will provide an invaluable resource for systematic chemical and genetic screens capable of reverting pathophysiological defects in physiology-relevant cellular models. This will facilitate the drug discovery and development for treating dystroglycanopathies.

## Experimental procedures

### Generation of human FKRP-iPSC lines

We obtained FKRP patient fibroblast lines from the MRC Centre for Neuromuscular Diseases Biobank. Approval for use of these cells has been in compliance with national guidelines regarding the use of biopsy tissue for research (REC reference 13/LO/1826; IRAS project ID: 141100). All patients or their legal guardians gave written informed consent. To generate patient-specific induced pluripotent stem cells (iPSCs), we implemented an efficient reprogramming technology based on a doxycycline-inducible system using six factors, OCT4, SOX2, KLF4, c-MYC, RARG and LRH1 (Wang et al., 2011). In vitro differentiation and analysis were carried out as described (Wang et al., 2011).

### Vector construction and CRISPR/Cas9 mediated genome editing

For targeted gene correction, we used Gibson Assembly (New England BioLabs) to construct the targeting donor vector. Briefly, the 1-kb left and right homology arms (LHA and RHA) were PCR-amplifed from parental *FKRP* fibroblast. The *FKRP* c.1364C>A (p.A455D) mutation on the LHA was simultaneously corrected with a modified primer (Table S2). The *piggyBac (PGK-puroΔtk)* selection cassette and vector backbone were PCR-amplified from the pMCS-AAT_PB-PGKpuroTK plasmid (Yusa et al., 2011). The four PCR fragments have 40-bp overlappings from end to end (primers are listed in Table S2) and joined together using Gibson Assembly Master Mix. The hCas9 plasmid was a gift from George Church (Addgene plasmid # 41815) (Mali et al., 2013). Short DNA fragments containing the *FKRP* sgRNA target sequence (Table S1) were generated by annealing two primers (Table S2), which create two overhangs ready for cloning into BsaI-digested p1261_U6_BsaI_gRNA_plasmid (kind gift of Sebastian Gerety, Wellcome Trust Sanger Institute). The expression of sgRNA is under the control of U6 promoter. The cell suspension was transferred to a cuvette and electroporated using Amaxa Nucleofection 2b device with program B16. For targeted gene mutation, the same strategy was used to construct the donor targeting vector carrying the *FKRP* c.1364C>A (p.A455D) mutation on LHA using primers in Table S2.

### Karyotyping analysis

As previously described (Agu et al., 2015), multiplex fluorescent in situ hybridization (M-FISH) karyotype analysis was performed on iPSC lines with slight modification. Briefly, prior to metaphase harvesting, iPSCs were grown in M15 medium (Knockout DMEM, 15% Fetal Bovine Serum, 1X glutamine-penicillin-streptomycin, 1X nonessential amino acids and 1 ng/ml human recombinant LIF) for 24 h and then treated with 10 μM Y-27632 dihydrochloride (Tocris) for 2–3 h.

### Neural induction and cortical differentiation from human iPSCs

As described (Yan et al., 2013), induction of NSCs from human iPSCs was carried out using Gibco Neural Induction Medium (Neurobasal medium and 2% Neural Induction Supplement) for 7days, followed by passaging and expansion in Neural Expansion Medium (49% Neurobasal medium, 49% Advanced DMDM/F-12 and 2% Neural Induction Supplement). Expanded NSCs were cryopreserved or differentiated into cortical neurons following the established protocol (Shi et al., 2012b). Briefly, tissue culture plates were pre-coated with 0.01% (w/v) poly-L-ornithin (Sigma, P4957) for 4 h, followed by 20 μg/ml of laminins (Sigma, L2020) for 4h. NSCs were seeded on the pre-coated plates (50,000/cm^2^) in Neural Expansion Medium with 5 μM Y-27632 dihydrochloride (Tocris) for 24 h. Subsequently, Neural Expansion Medium was replaced with Neural Maintenance Medium, containing 50% DMEM/F-12 (Gibco), 50% Neurobasal medium (Gibco), 0.5X N2 supplement (Gibco), 0.5X B27 supplement (Gibco), 1.5 mM GlutaMAX-I (Gibco), 0.5X Penicillinstreptomycin (Gibco), 2.5 μg/ml Insulin (Sigma), 0.05 mM of 2-Mecaptoethanol (Gibco), 0.5% v/v of Non-Essential Amino Acid (Gibco), 20 ng/ml BDNF (Peprotech) and 20 ng/ml GDNF (Peprotech). When reaching 90% confluence (approximately within 3-4days), cells were re-seeded (60,000/cm^2^) in Neural Maintenance Medium (with 5 μM Y-27632 dihydrochloride for 24 h), followed by media change every other day. In vitro cortigenesis occurs in the following weeks. Cultured cells were then harvested or fixed at specific time points for further analysis.

### Immunocytochemistry

Cells were fixed with 4% PFA for 15 mins at room temperature (RT). Prior to immunocytochemistry, cells were permeabilized with 0.1% Triton/phosphate-buffered saline (PBS) for 15min at RT and blocked with either 10% goat serum/PBS or 10% horse serum/PBS. Primary antibodies used were: OCT4 (1:100; Santa Cruz, sc-5279), NANOG (1:100; abcam, AB80892), Tra-1-60 (1:100; Santa Cruz, sc-21705), SSEA4 (1:100; BD Bioscience, 560796), α-fetoprotein (1:150; R&D Systems, MAB1368), α-smooth muscle actin (1:150; R&D Systems, MAB1420), Sox1 (1:100; R&D Systems, AF3369), Sox2 (1:100; R&D Systems, 245610), Nestin (1:500; abcam, ab22035), Ki67 (1:100; BD Pharmingen, 550609), OTX2 (1:250; Millipore, AB9566), Pax6 (1:300; Biolegend, 901301), vimentin (1:100; abcam, ab28028), vGLUT1 (1:2000; Synaptic systems, 135303) and β-III tubulin (Tuj1) (1:150; R&D System, MAB1195). All primary antibodies were diluted in blocking solution and incubated overnight at 4°C, followed by washing in PBS 3 times for 15 mins. Subsequently, appropriate Alexa Fluor 488 or 564 conjugated secondary antibodies were incubated for 1 h at RT. For nuclei staining, samples were incubated with DAPI (Sigma-Aldrich) for 5 mins and washed in PBS briefly. Finally, images were captured and analysed using IN Cell 2200.

### Immunoblot analysis

Cells were harvested at specific time points and proteins were extracted in RIPA buffer consisting of 50 mM Tris-HCl (pH 7.5), 150 mM NaCl, 1 mM EDTA, 1% Triton X-100, 1% SDS and 1 mM azide plus a cocktail of protease inhibitors (Roche). A total of 30-60 μg of soluble protein was resolved using NuPage Novex 4-12% Bis-Tris protein gels (Invitrogen) and then electrophoretically transferred to polyvinylidene difluoride (PVDF) membrane (Millipore). The PVDF membrane was blocked in 3% bovine serum albumin in TBST (Tris-buffered saline with 0.1% Tween) and probed with primary antibodies to glycosylated α-dystroglycan (IIH6 1:500; kind gift from Kevin Campbell) and β-dystroglycan (1:100; Leica Biosystems, NCL-b-DG), followed by washing with TBST and incubation with horseradish peroxidase (HRP) conjugated anti-mouse IgM (Millipore) and HRP conjugated anti-mouse IgG (Jackson laboratories), respectively. Note that anti-beta actin antibody (1:1000; abcam, ab8226) used in this study does not cross-react with skeletal muscle actin. SuperSignal West Pico Chemiluminescent Substrate (Thermo Fisher) was applied to PVDF membranes and signals were visualized using a Bio-Rad ChemiDoc MP imaging system.

### Laminin overlay analysis

Protein samples were extracted, resolved and transferred to PVDF membranes as described above. The PVDF membrane was then blocked with 5% skimmed milk in laminin binding buffer (LBB), containing 10 mM triethanolamine, 140 mM NaCl, 1 mM MgCl_2_, 1 mM CaCl_2_ (pH adjusted to 7.6), followed by incubation with laminins from Engelbreth-Holm-Swarm murine sarcoma basement membrane (Sigma, L2020) in LBB with a final concentration of 5 μg/ml at 4^o^C overnight. The PVDF membrane was then incubated with the pan-laminin antibody (1:1000; Sigma, L9393), washed with 1x LBB with 0.1% Tween and incubated with HRP conjugated anti-rabbit IgG (Jackson laboratories). Signal detection using chemiluminescence substrate was performed as above.

## Author Contribution

B.L., J.K., D.R., and E.K. carried out the experimental work, analyzed and interpreted the data. S.G, B.F, and F.Y. performed and analyzed the karyotyping experiments. D.R., D.S., P.L., and F.M. contributed to the conception and design of the project and edited the manuscript. Y.-Y.L. designed the project, performed some of the experiments, analyzed and interpreted the data and supervised the research. J.K., and Y.-Y.L. co-wrote the manuscript.

## Acknowledgement

We thank Dr Wei Wang (Wellcome Trust Sanger Institute) for his critical advice on the generation of human iPSCs. We are indebted to Dr Kosuke Yusa (Wellcome Trust Sanger Institute) for valuable discussion about genome editing and providing plasmids. We would like to thank Dr Susan Brown (Royal Veterinary College) for providing wildtype mouse muscle and brain lysates and critical reading of the manuscript. We thank Dr Luke Gammon (Blizard Institute Screening Core Facility) and Amaia Paredes Redondo (Blizard Institute) for their kind assistance in some experiments. F.M. is supported by the National Institute for Health Research Biomedical Research Centre at Great Ormond Street Hospital for Children NHS Foundation Trust and University College London. The support of the Muscular Dystrophy UK to the Dubowitz Neuromuscular Centre and of the MRC Neuromuscular Centre Biobank is also gratefully acknowledged. This work was supported in part by Royal Society Research Grant (RG130417) and Newlife Research grant (SG/14-15/14) to Y.-Y.L.

